# A Lung Cancer Mouse Model Database

**DOI:** 10.1101/2024.02.28.582577

**Authors:** Ling Cai, Ying Gao, Ralph J. DeBerardinis, George Acquaah-Mensah, Vassilis Aidinis, Jennifer E. Beane, Shyam Biswal, Ting Chen, Carla P. Concepcion-Crisol, Barbara M. Grüner, Deshui Jia, Robert Jones, Jonathan M. Kurie, Min Gyu Lee, Per Lindahl, Yonathan Lissanu, Maria Corina Lorz Lopez, Rosanna Martinelli, Pawel K. Mazur, Sarah A. Mazzilli, Shinji Mii, Herwig Moll, Roger Moorehead, Edward E. Morrisey, Sheng Rong Ng, Matthew G. Oser, Arun R. Pandiri, Charles A. Powell, Giorgio Ramadori, Mirentxu Santos Lafuente, Eric Snyder, Rocio Sotillo, Kang-Yi Su, Tetsuro Taki, Kekoa Taparra, Yifeng Xia, Ed van Veen, Monte M. Winslow, Guanghua Xiao, Charles M. Rudin, Trudy G. Oliver, Yang Xie, John D. Minna

## Abstract

Lung cancer, the leading cause of cancer mortality, exhibits diverse histological subtypes and genetic complexities. Numerous preclinical mouse models have been developed to study lung cancer, but data from these models are disparate, siloed, and difficult to compare in a centralized fashion. Here we established the Lung Cancer Mouse Model Database (LCMMDB), an extensive repository of 1,354 samples from 77 transcriptomic datasets covering 974 samples from genetically engineered mouse models (GEMMs), 368 samples from carcinogen-induced models, and 12 samples from a spontaneous model. Meticulous curation and collaboration with data depositors have produced a robust and comprehensive database, enhancing the fidelity of the genetic landscape it depicts. The LCMMDB aligns 859 tumors from GEMMs with human lung cancer mutations, enabling comparative analysis and revealing a pressing need to broaden the diversity of genetic aberrations modeled in GEMMs. Accompanying this resource, we developed a web application that offers researchers intuitive tools for in-depth gene expression analysis. With standardized reprocessing of gene expression data, the LCMMDB serves as a powerful platform for cross-study comparison and lays the groundwork for future research, aiming to bridge the gap between mouse models and human lung cancer for improved translational relevance.

## Introduction

Lung cancer remains the most common cause of cancer-related mortality globally, with its complexity reflected in diverse histological subtypes—such as adenocarcinoma (ADC), squamous cell carcinoma (SQCC), large cell carcinoma (LCC), and small cell lung cancer (SCLC)—each harboring distinct genetic alterations that drive tumor biology, which in some cases dictates therapeutic vulnerabilities. To decipher the complexities of tumor biology, high-throughput molecular profiling of patient-derived tumors has been extensively employed [1-7].

Preclinical models of lung cancer are essential tools for researchers to understand cancer biology and develop therapeutic strategies through experimentation. There has been a concerted effort to aggregate data from patient-derived cell lines [8, 9] and patient-derived xenografts (PDXs) [10]. While lung cancer autochthonous animal models, primarily based on mice, represent a separate but significant line of research, they often lack unified characterization due to independent development across various laboratories.

To address this gap, we conducted a comprehensive review of transcriptomic databases, specifically GEO and ArrayExpress, collected transcriptomic data from lung cancer mouse models, and standardized associated sample and oncogenotype information. We actively engaged with data depositors to refine our curation process and incorporate their insights. These efforts have culminated in the creation of the Lung Cancer Mouse Model Database (LCMMDB). This resource serves as a centralized platform for the research community, providing access to a comprehensive collection of genetically engineered and chemically induced mouse models of lung cancer. We also developed a user-friendly web application populated from this database, offering researchers intuitive tools for dynamic data exploration and sophisticated analysis.

## Methods

### Dataset screening

We performed searches with keywords “lung cancer” refined to species “Mus Musculus” and data type restricted to gene expression profiling by array or high-throughput sequencing in the Gene Expression Omnibus (GEO) and ArrayExpress. Each of these studies was manually inspected to identify gene expression data generated from autochthonous models, including genetically engineered mouse models (GEMMs), chemically induced mouse models, and spontaneous models of lung cancer. R package GEOquery [11] was used to download author-processed gene expression data and sample annotation data from GEO.

### LCMMDB data organization and curation process

LCMMDB organizes data into three primary tables, for datasets, samples, and genotypes. The dataset table contains data accession IDs, platform IDs, model types, study titles, publication PMIDs, PMC IDs, and the contact information of data depositors. The sample table contains accession IDs, sample names, types, treatments, strains, sex, age, genotype, histological classification, primary/metastasis status, sources of Affymetrix data, Short Read Archive (SRA) IDs, and growth protocols. The genotype table was designed to record details of model genetic manipulations. It contains multiple rows for each genotype to specify genes involved, genetic constructs, zygosity, type of genetic modification (e.g., overexpression, knockout), method of genetic manipulation, induction methods, induction systems, promoters used, cell of origin, and additional notes that may provide context or clarifications. This information is further organized to generate both standardized and simplified genotypes, concisely indicating the genetic manipulations and induction methods employed in each model.

For data curation, we gathered details from database annotations and carefully reviewed the original publications to extract the necessary information. We standardized terms to ensure consistency across the data. For instance, we categorized sample types into four distinct groups: “bulk tissue,” “microdissected,” “CD45 depleted,” and “sorted cancer cells”. We also include data fields for the original curation to preserve the intricacies of the source dataset. For example, while we simplified the primary/metastasis tumor status to “primary” and “metastasis” for consistency, we kept specific details like “liver metastasis” in the “primary/metastasis original” field to capture the full depth of the original classifications.

In harmonizing the histology data, we recognized the continuum that exists between mouse tumor classifications of adenoma and adenocarcinoma (ADC). For example, in the LSL-K-ras^G12D^ model, tumors can progress from adenoma to adenocarcinoma between 6-16 weeks post-infection [12]. However, not all studies explicitly differentiate between adenoma and adenocarcinoma. Additionally, multiple clonal tumors may present within the same sample, where some may classify as adenomas and others as adenocarcinomas. To address this, we carefully reviewed original publications and annotations, assigning the most accurate histology annotation to the “histology.original” field. For cases with clear distinctions, we labeled them as either “Adenoma” or “ADC.” For those with ambiguous classifications, we used “Adenoma/ADC.” Consequently, in the “histology” field, we grouped these classifications together under “Adenoma/ADC” to maintain consistency and clarity across the dataset.

### Gene expression reprocessing

Affymetrix raw data were downloaded from GEO and grouped by platform. For each platform, we downloaded v25 of the gene-level customized Chip Definition Files (CDFs) from the Molecular & Behavioral Neuroscience Institute (MBNI) repository (http://brainarray.mbni.med.umich.edu/Brainarray/Database/CustomCDF/25.0.0/ensg.asp) at the University of Michigan [13], to reprocess the data with the most up-to-date and specific gene annotations. The CEL files were batch-read with the specified platform package and normalized using the Robust Multi-array Average (RMA) method via the oligo package, yielding an expression set (eset) from which gene expression matrices were extracted. Entrez IDs were converted to gene symbols based on the NCBI Entrez mapping file.

RNA-seq fastq files were downloaded from Sequence Read Archive (SRA) through SRA-Toolkit. Paired-end reads were concatenated to be processed as single-end reads. Reads were trimmed to remove adapters and low-quality sequences and subsequently aligned to mouse reference GRCh38m by hisat2 [14]. Gene expression was quantified using FeatureCounts [15] and GENOCODE [16]. We retained genes with non-zero values in more than 10% of samples, normalized their counts to library sizes, and computed log-transformed counts per million (logCPM) values for downstream analyses.

### AACR GENIE data analysis

AACR GENIE data [17] (Version 15.0-public) was downloaded from SAGE BIONETWORKS on 03/25/2024 through R package “synapserutils”[18] with Synapse ID “syn7222066”. We used mutation status from “data_mutations_extended.txt”, amplification status (value of 2) and deletion status (value of -2) from “data_CNA.txt” and structural variation (sv) status from “data_sv.txt” to determine genetic aberrations. Lung cancer patient samples were selected from “data_clinical_sample.txt”. Cumulative counts of genetic aberration events are summarized at the sample level (note that some patients could have multiple samples in the dataset).

### Web application construction

Statistical software R was used for analyses and web application construction [19]. The web application https://lccl.shinyapps.io/LCMMDB/ is a shiny app deployed at the shinyapps.io servers. It is implemented through the following R packages: ‘data.table’, ‘reshape2’, ‘digest’, ‘stringr’, ‘dplyr’, ‘tidyverse’, ‘Hmisc’, ‘ggplot2’, ‘RColorBrewer’, ‘ggpubr’, ‘patchwork’, ‘shinyjs’, ‘shinycssloaders’, ‘bslib’, ‘htmltools’, ‘shinyWidgets’, ‘shinyTree’, ‘DT’, and ‘plotly’.

## Results

### Construction of the Lung Cancer Mouse Model Database (LCMMDB)

An exhaustive search in the GEO and ArrayExpress identified nearly 500 candidate lung cancer autochthonous mouse model datasets. Each of these studies was manually inspected to identify transcriptomic data generated from GEMMs, chemically induced tumors, or spontaneously formed tumors. Additionally, we included control lung samples and those exposed to carcinogenic treatments while excluding mouse cell lines and allografts into syngeneic recipients to ensure specificity to our research focus. We removed two datasets due to data redundancy from the reprocessed data (**Figure S1**). Our current data collection includes 77 datasets from 71 unique studies, comprised of 1,354 samples (**Table 1** and **Figure 1a**).

**Table 1.**
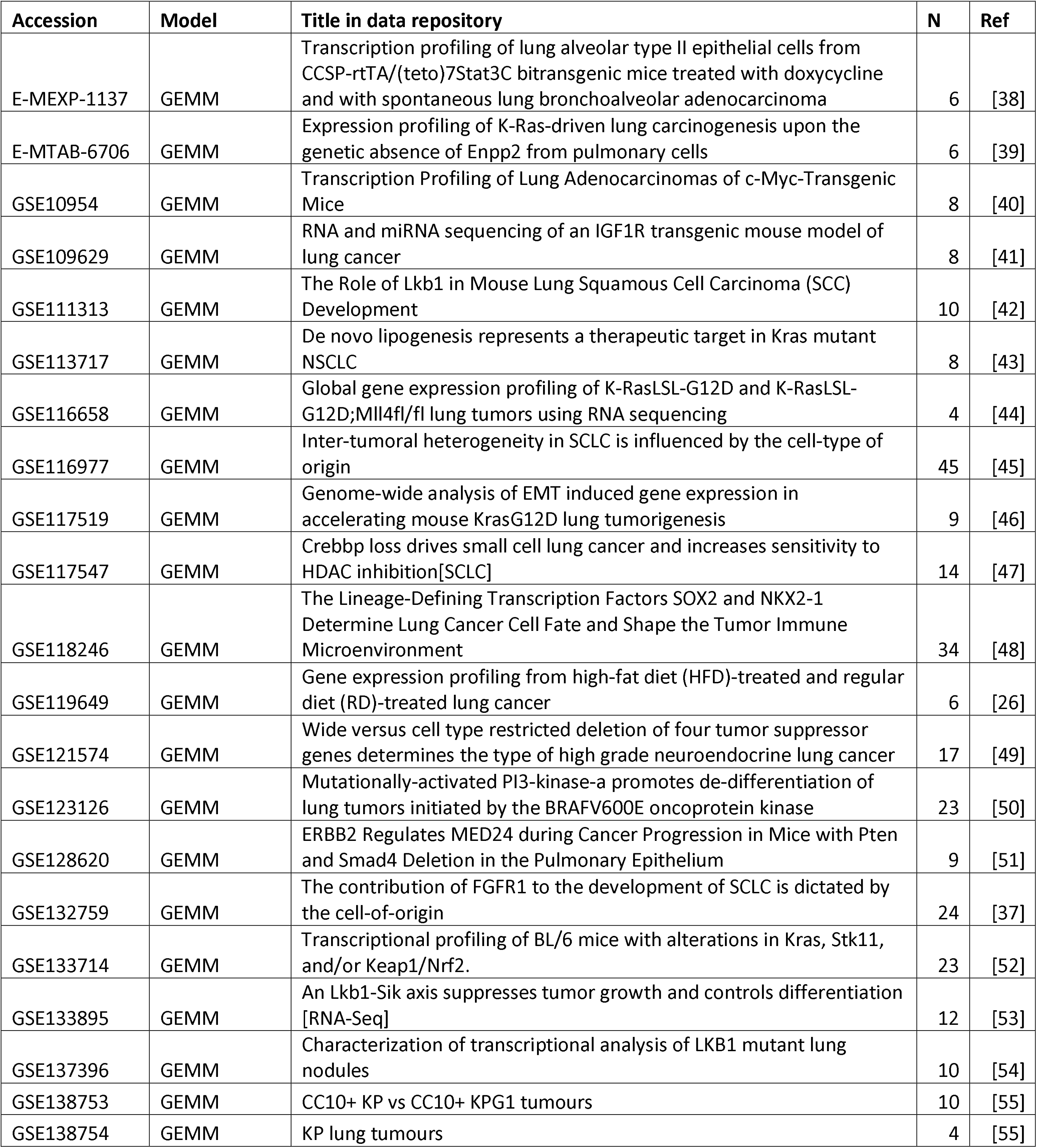
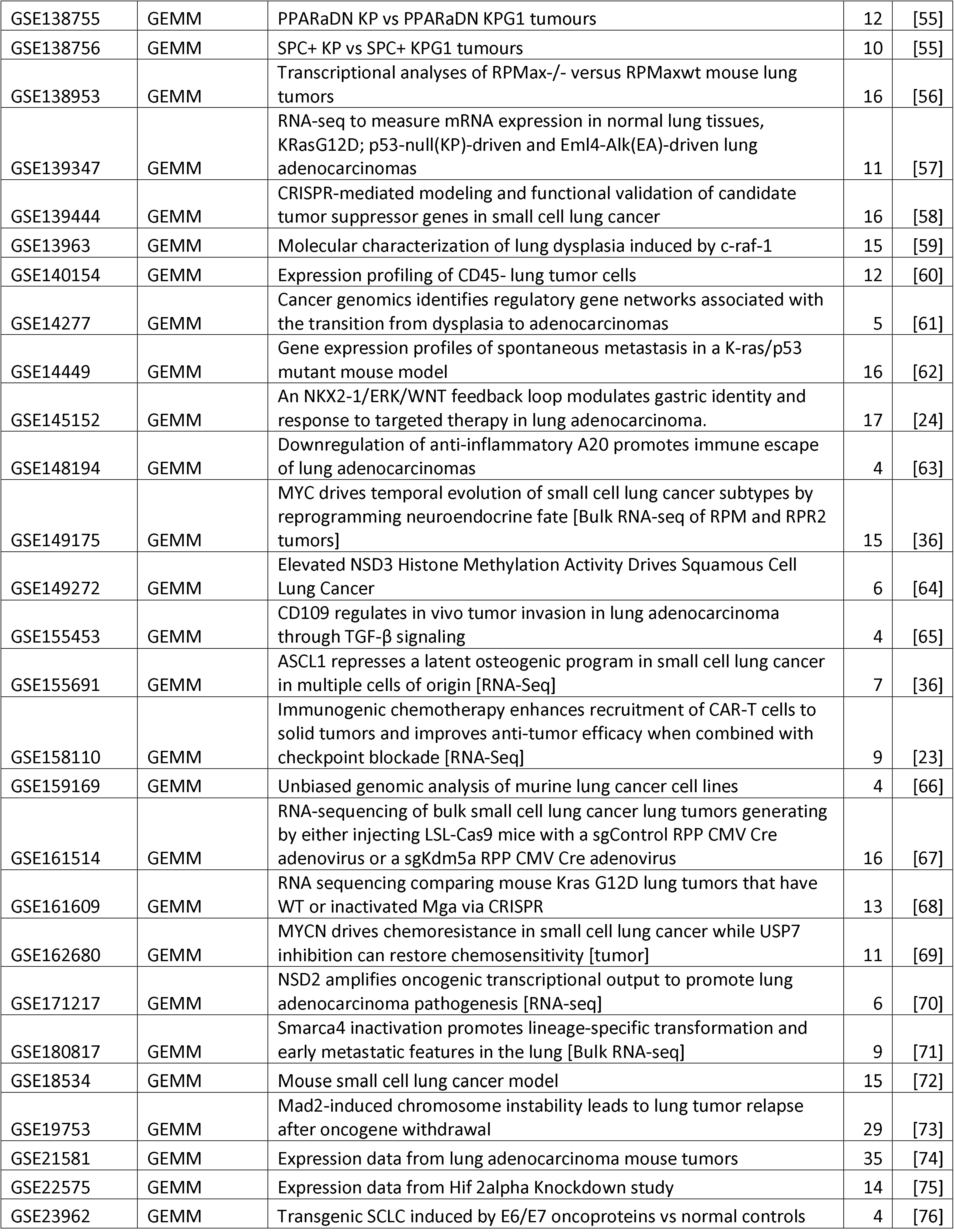
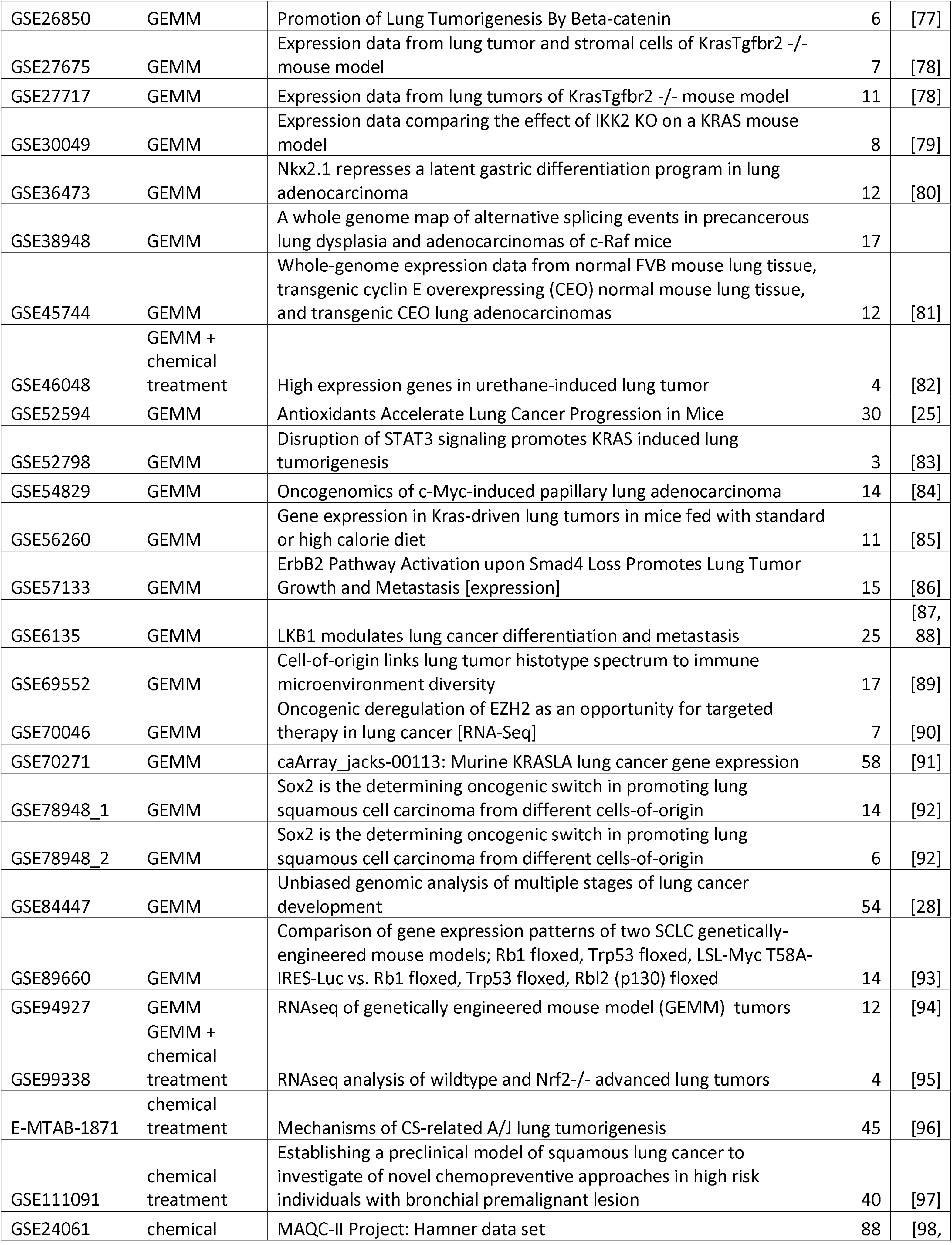
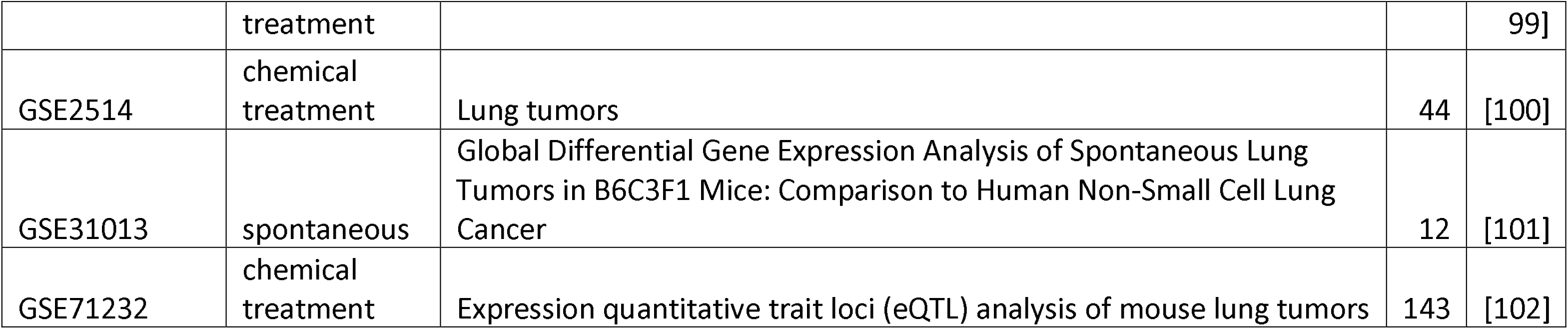
Current collection of LCMMDB datasets.

**Figure 1.**
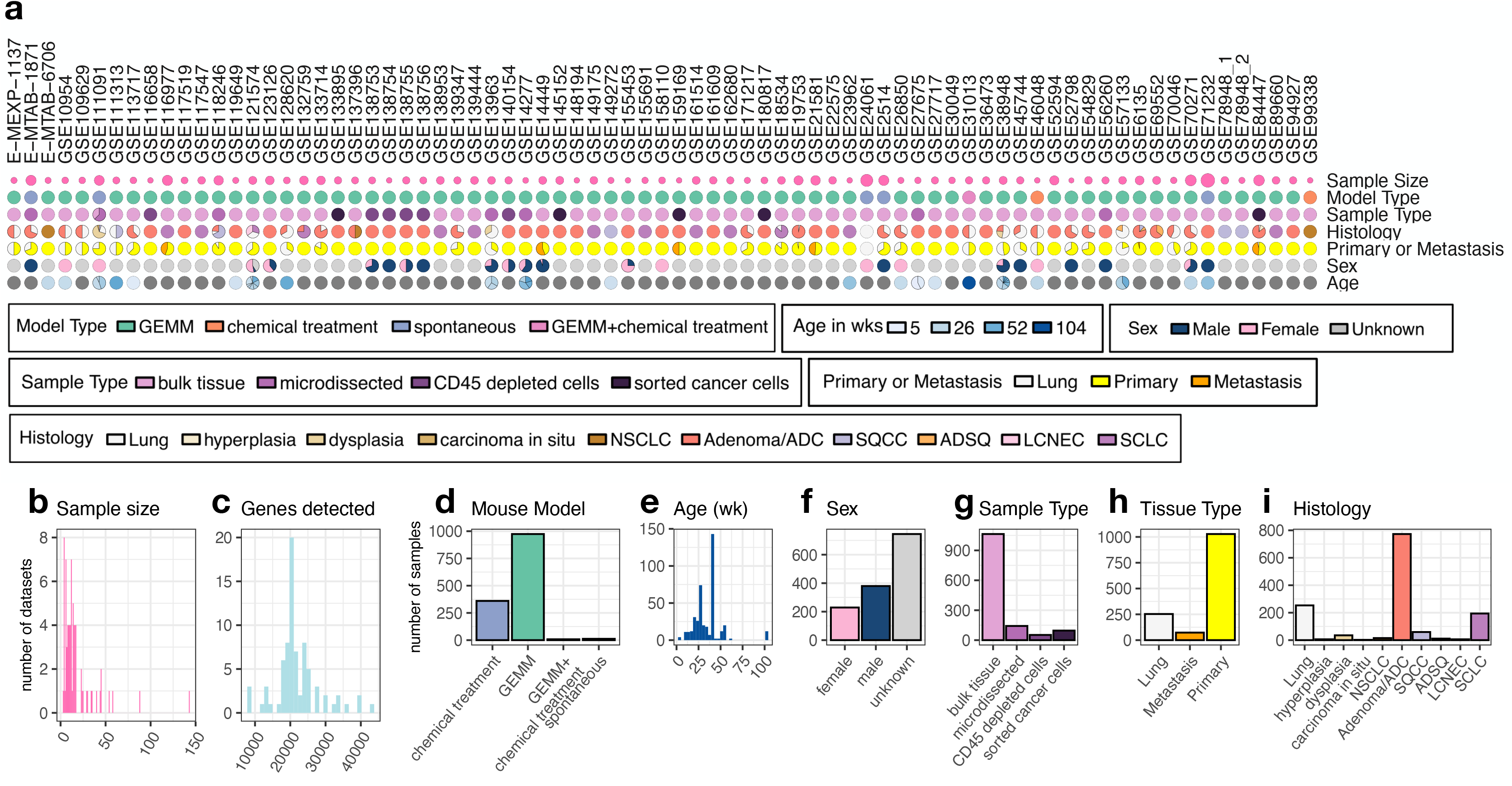
Overview of Sample Characteristics and Distribution in LCMMDB. **a**. Characteristics of individual datasets by pie charts. Each column represents a dataset, and each row corresponds to a specific attribute, with color-coding denoting the category. Attributes include Model Type, Age, Sex, Sample Type, Histology, and Primary or Metastasis status. Dark gray color denotes missing data. **b-c**. Sample size (**b**) and gene feature number (**c**) distribution across all datasets by bar plots. **d**-**h**. Distribution of samples by Model Type (**d**), Age (**e**), Sex (**f**), Sample Type (**g**), Tissue Type (**h**) and Histology (**i**). Note that “Lung” under Tissue Type or Histology can include normal wildtype lungs but also chemical treated lungs from toxicology studies, or genetically modified non-wildtype lungs.

After a thorough data harmonization process, we contacted the data depositors and shared the curated data specific to their studies along with our data schema, soliciting their verification, rectifications, or any insights they could offer. 89% of the data depositors responded to our request to confirm our data curation. More than half of these contributors provided valuable corrections and insights, with some recommending additional datasets for future inclusion (detailed in **Table S1**). The database was updated accordingly, integrating the depositors’ revised data and constructive feedback.

Our analysis revealed a general trend towards small sample sizes across the datasets, with a median of 12 samples, ranging from 3 to 143 (**Figure 1b**). The median number of detected genes per study is 20,942, with older microarray platforms reporting fewer genes (**Figure 1c**). The majority of the samples originated from 71 GEMM datasets, including 856 cancerous samples and 118 lung samples. Complementary to these, seven studies offered 368 samples generated from carcinogen-exposed models, including 239 cancerous samples and 129 lung samples. One unique dataset provided insight into 6 spontaneous tumors and 6 lung samples in mice aged two years (**Figure 1d**). Age details were available for 407 samples (30%), with a median age of 34 weeks (**Figure 1e**). Sex annotations were available for 609 samples (45%), encompassing 11 datasets with mixed sexes, six with exclusively female mice, and eight with exclusively male mice (**Figure 1f**).

The sample types were predominantly from bulk tissue or microdissected specimens. A subset of 146 samples from 11 datasets underwent techniques such as CD45-depletion or fluorescence-based cancer cell sorting to reduce tumor microenvironmental contributions (**Figure 1g**). Within the 1,101 curated tumor samples, 73 were identified as metastatic, with 53 metastases arising from ADC primary tumors and 20 from SCLC (**Figure 1h**).

We curated 197 adenomas, 337 ADCs, and 236 cases classified as both based on authors’ reports and literature reviews. Due to the overlapping lineage relationship of adenoma and ADC classifications, we opted to aggregate these under the “Adenoma/ADC” category for standardized histological classification. This aggregation highlighted a dataset composition with 73% Adenoma/ADC, 18.3% SCLC, and only 5.6% SQCC, indicating an underrepresentation of SQCC when contrasted with its prevalence in human lung cancer (**Figure 1i**).

### The genetic landscape of GEMMs and comparison with patient mutation spectrum

859 precancerous lesions (hyperplasia, dysplasia, carcinoma *in situ*) and tumor samples in the LCMMDB collection were developed from GEMMs. We curated genotype tables to record the involved genes, allele zygosity, genetic modifications, manipulative techniques, induction methods, and cells of origin. We illustrate the genetic alteration landscape, involving either single or combined manipulations of 54 genes in these GEMM samples in **Figure 2a**. These include six human genes (*EGFR, IGF1R, EZH2, MYCN, CCNE1*, and *SNAI1*) and two viral genes (HPV *E6* and *E7*) introduced to the GEMMs. We compiled standardized genotypes and simplified them to harmonize genotype curation and identified a total of 122 unique standardized genotypes.

**Figure 2.**
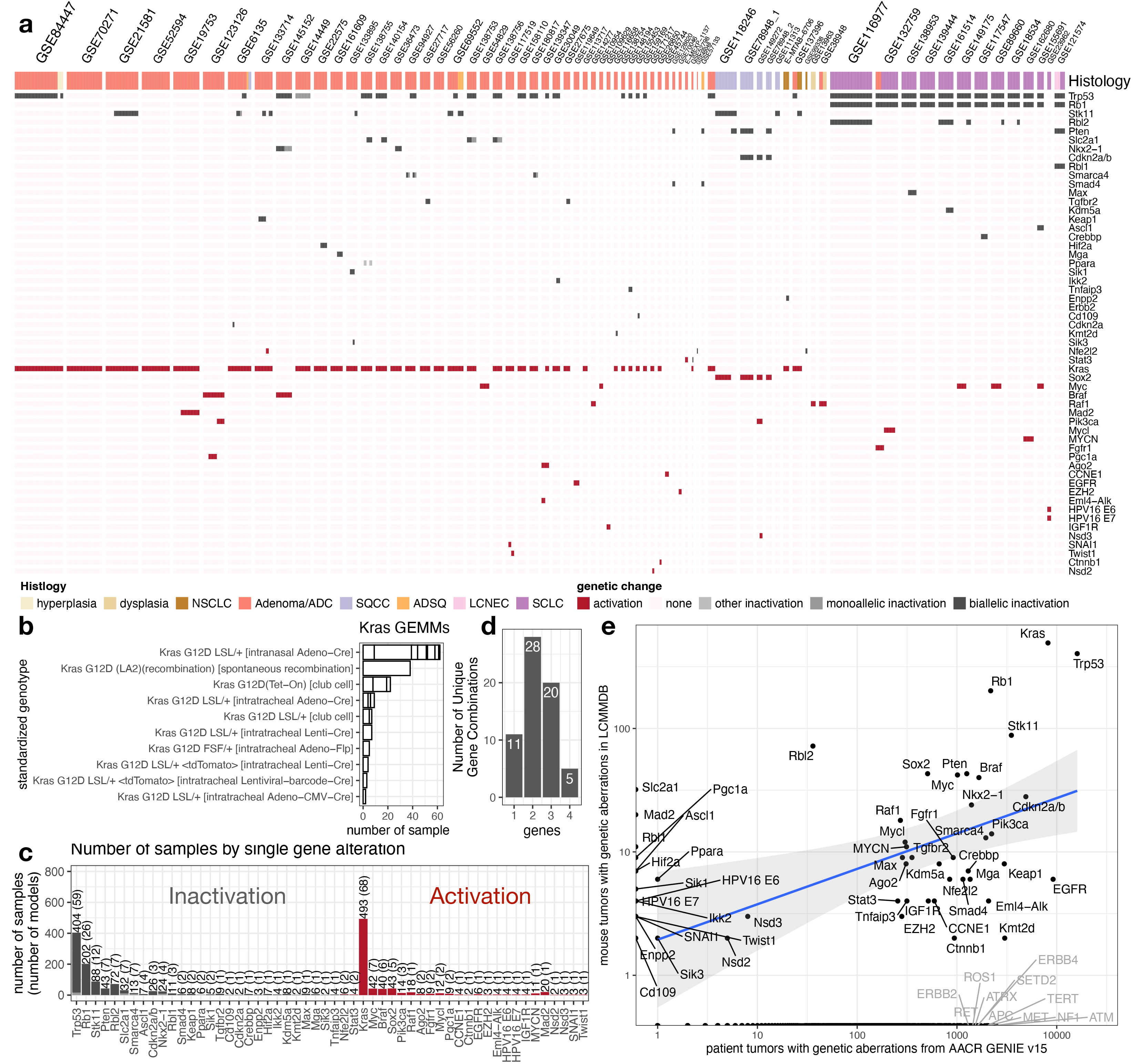
Summary of GEMM genotypes in LCMMDB. **a**. Landscape of genetic modifications in LCMMDB GEMM tumors by dataset and histology. **b**. Sample count by standardized genotype in GEMM tumors with Kras mutation alone. Small boxes within the bars represent samples from different datasets. **c**. Sample count in GEMM tumors by single gene alteration. Y-axis labels indicate the total number of tumors with the specific gene altered and the number of unique standardized genotypes with the specified gene altered in parentheses. **d**. Count of GEMMs by the number of altered genes. Bars represent the number of GEMMs with one to four manipulated genes, irrespective of manipulation method or mutation. **e**. GEMM tumor alterations in LCMMDB vs. human lung cancer genetic aberrations in the AACR GENIE v15 database by gene. Selected human oncogene and tumor suppressor genes not represented in LCMMDB are highlighted in gray.

Considering lesions/tumors arising from *Kras* manipulation alone, for example, 10 distinct standardized genotypes were identified, which vary in genetic constructs and induction methods (**Figure 2b**). Remarkably, all these genotypes harbor the G12D mutation, representing only ∼15% of *KRAS* mutations in non-small cell lung cancer (NSCLC) patients [20]. This disparity underscores the broader issue of limited genetic variation in GEMM tumors compared to human cancers, which is also exemplified by *Trp53* mutations. Beyond simply inactivating p53, mutations in this gene are known to confer additional gain-of-function properties [21]. However, in our current LCMMDB database, out of 404 GEMM tumors with *Trp53* manipulation, only 16 samples originate from a single study using a *Trp53*^R172H^ mutant model, with the remainder predominantly involving knockouts or knockdowns.

The gene-centric distribution of genetic alterations in the tumor samples is detailed in **Figure 2c**. Twenty-eight genes are predominantly activated while 24 are primarily inactivated. Two genes, *Nfe2l2* and *Stat3*, were subject to both activation and inactivation studies within the GEMMs. This figure also denotes the number of standardized genotypes associated with each gene, represented in parentheses next to the total sample count. Notably, 29 of the 54 genes were exclusive to a single model. When considering the unique gene combinations, dual-gene manipulations emerged as the most common scenario, presented in 28 distinct instances. In contrast, manipulations of 11 different single genes were adequate to generate GEMM tumors (**Figure 2d**). Only five models contained alterations in 4 genes (**Figure 2d**), likely reflecting the inherent challenges associated with the time and expense required to generate mice with quadruple-modified alleles.

We next performed a comparative analysis of the frequency of genetic alterations in mouse lung cancer GEMM-derived tumors with those identified in human lung cancers (**Figure 2e**), as recorded in the AACR GENIE v15 database, based on clinical sequencing data from real-world patient populations [17]. Mutations in *TP53* and *KRAS*, among the most prevalent mutations in human lung cancer, are adequately represented in the GEMM tumors. The observed positive correlation in gene alteration frequencies between mouse tumors and patient tumors suggests that GEMMs frequently incorporate genes commonly mutated in human lung cancer. However, our review indicates that some genes implicated in human lung cancers are understudied within the GEMM framework. For instance, the *Eml4-Alk* translocation and *Kmt2d* inactivation have each been characterized in only one study in our database. Moreover, pivotal oncogenes such as *ROS1, MET, RET, ERBB4*, and critical tumor suppressor genes like *NF1, ATM*, and *APC* are currently absent from the LCMMDB (**Figure 2e**). **Figure S2** details the frequency and types of genetic alterations for the top 100 genes most frequently altered in lung cancer patients according to the AACR GENIE data, with an emphasis on 78 genes that are not yet included in the LCMMDB. These findings underscore the need to broaden the scope of lung cancer GEMM development and characterization to cover a more extensive array of genetic drivers of the disease.

### Harmonization of gene expression data

To address the limited sample size within individual datasets, we acquired raw data where possible and reprocessed them through standardized pipelines by platform, each with the latest probe and gene annotations. This standardization effort enabled us to make reprocessed data available for 85% of the samples (**Figure 3a**). Notably, approximately half of these samples (n = 563) are derived from RNA-seq and encompass 38 distinct datasets (**Figure 3b**). Principal component analysis (PCA) conducted on the top 1000 variable genes from the reprocessed RNA-seq data revealed that the first two principal components (PCs) capture 62% of the total variance, indicating a strong structuring of the data (**Figure 3c**). Despite potential batch effects, the PCA demonstrates that different datasets exhibit substantial overlap (**Figure 3d**), with clear distinctions observed between SCLC and NSCLC samples along PC1, and between primary and metastatic samples along PC2 (**Figures 3e** and **3f**, respectively). For microarray datasets, we executed a parallel processing strategy on data from the Mouse430_2 platform—the most represented microarray platform with 283 samples across 15 datasets—and noted a comparable success in data integration (**Figure S3**). Although batches from various experimental conditions, sample types, and biological differences such as mouse age, sex, and strain may still be present, our reprocessing method appears to have effectively consolidated the datasets, thereby facilitating cross-dataset comparisons and potentially uncovering broader trends within the merged data.

**Figure 3.**
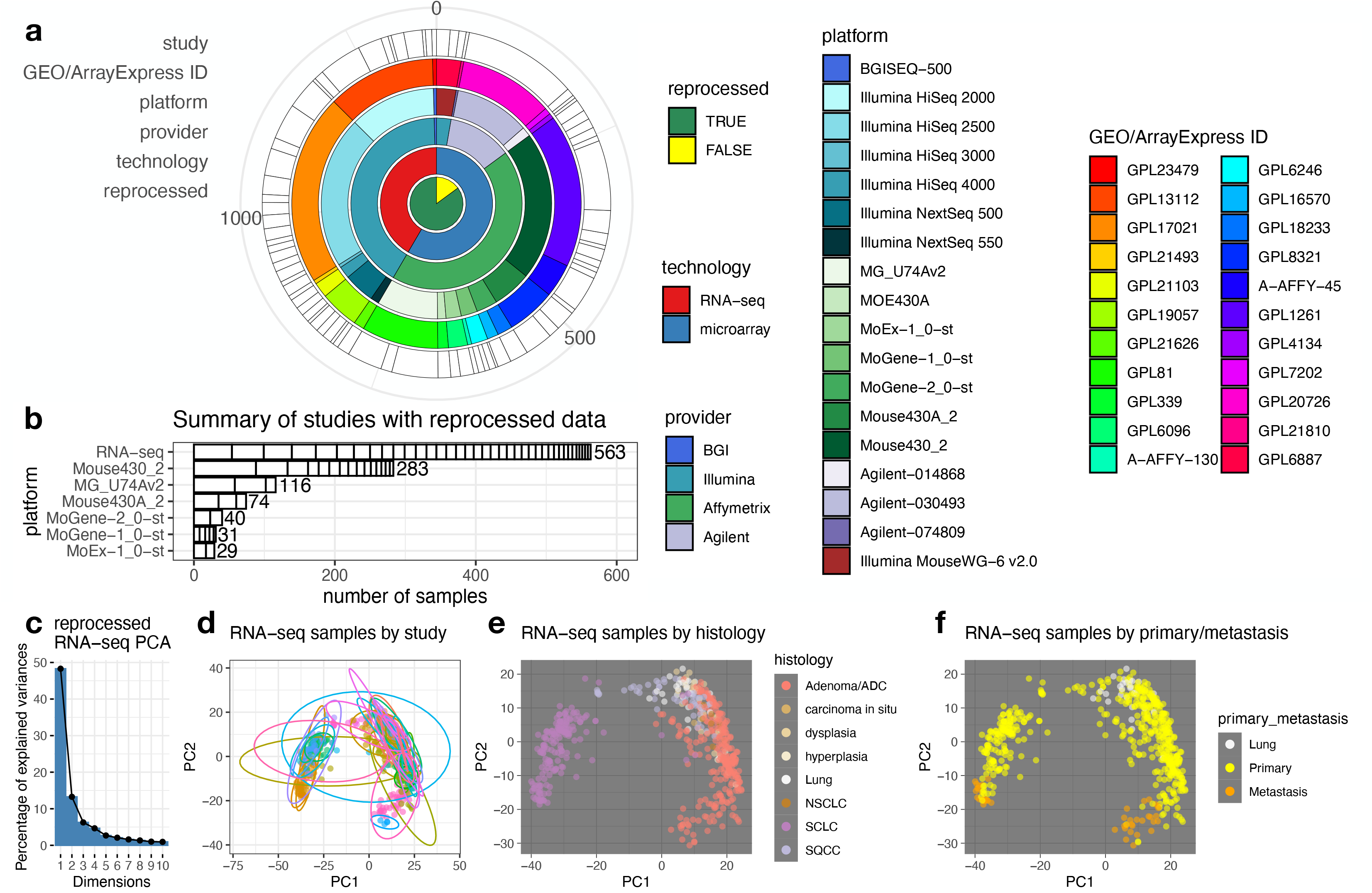
Data reprocessing by platform. **a**. Hierarchical relationship of technology and platforms. 85% (1152 samples) of the LCMMDB gene expression data was reprocessed. **b**. Platforms with multiple studies reprocesed through standardized workflow. Each box within the bars represents a single dataset. **c**. In principal component analysis using 1000 most variable genes from reprocessed RNA-seq data, the top two principal components accounts for 62% of the total variance. **d**-**f**. Distribution of 563 RNA-seq samples by source dataset (**d**), histology (**e**), and primary/metastasis status (**f**).

### A user-friendly web application for LCMMDB

To facilitate the exploration and analysis of the LCMMDB data, we constructed a web application that can be accessed at https://lccl.shinyapps.io/LCMMDB/. This application is structured into two primary sections: a data review panel and an analysis panel. Within the data review panel, the “Overview” tab presents graphical summaries of the LCMMDB, while the “Studies,” “Samples,” and “GEMMs” tabs allow users to navigate and refine detailed data tables. These tables correspond to **Supplementary Tables 2-5** in this manuscript. Specifically, the “GEMMs” tab displays a table where genetic alterations are recorded with one gene per line. Each genotype within a study is distinctively highlighted to ensure clear visual separation. Users can customize their view, choosing which columns to display and applying filters to refine row entries—such as querying specific gene combinations, with an illustrative example in **Figure S4**.

The analysis panel offers users an interactive environment to delve deeper into the gene expression profiles across multiple datasets. The “Depositor-processed” data option allows researchers to analyze the expression data as originally submitted, maintaining consistency within datasets and enabling reliable within-dataset comparisons. The results are visualized as a series of dot plots, ranked by the statistical significance of expression differences determined by one-way ANOVA. The “Merged by platform” data option allows users to examine the reprocessed data by platform, leveraging the harmonized datasets to discern patterns and insights across different studies.

### Comparisons in individual datasets

We offer three options for gene expression comparison using depositor-processed data. The first, which compares expression by genotype and/or treatment, provides the broadest dataset range and includes versatile sample filtering capabilities. Users can refine the analysis parameters by utilizing the available filters within the dropdown menu, tailoring the analysis to their specific research interests (**Figure S5**). As exemplified in **Figure 4**, where the top 6 of 44 datasets qualified from the specified criteria are shown, we identified several genetic and treatment conditions that induced the most prominent *Cd274* (PD-L1) expression changes in NSCLC bulk tissue samples. This is particularly notable in models with *Stk11* (also known as *Lkb1*) knockout, where *Cd274* expression is markedly downregulated, corroborating clinical findings that *STK11* mutations are significantly enriched among PD-L1-negative lung tumors [22]. On the other hand, treatment of oxaliplatin and cyclophosphamide (Ox/Cy)[23], known to induce immunogenic cell death increased the expression of *Cd274*.

**Figure 4:**
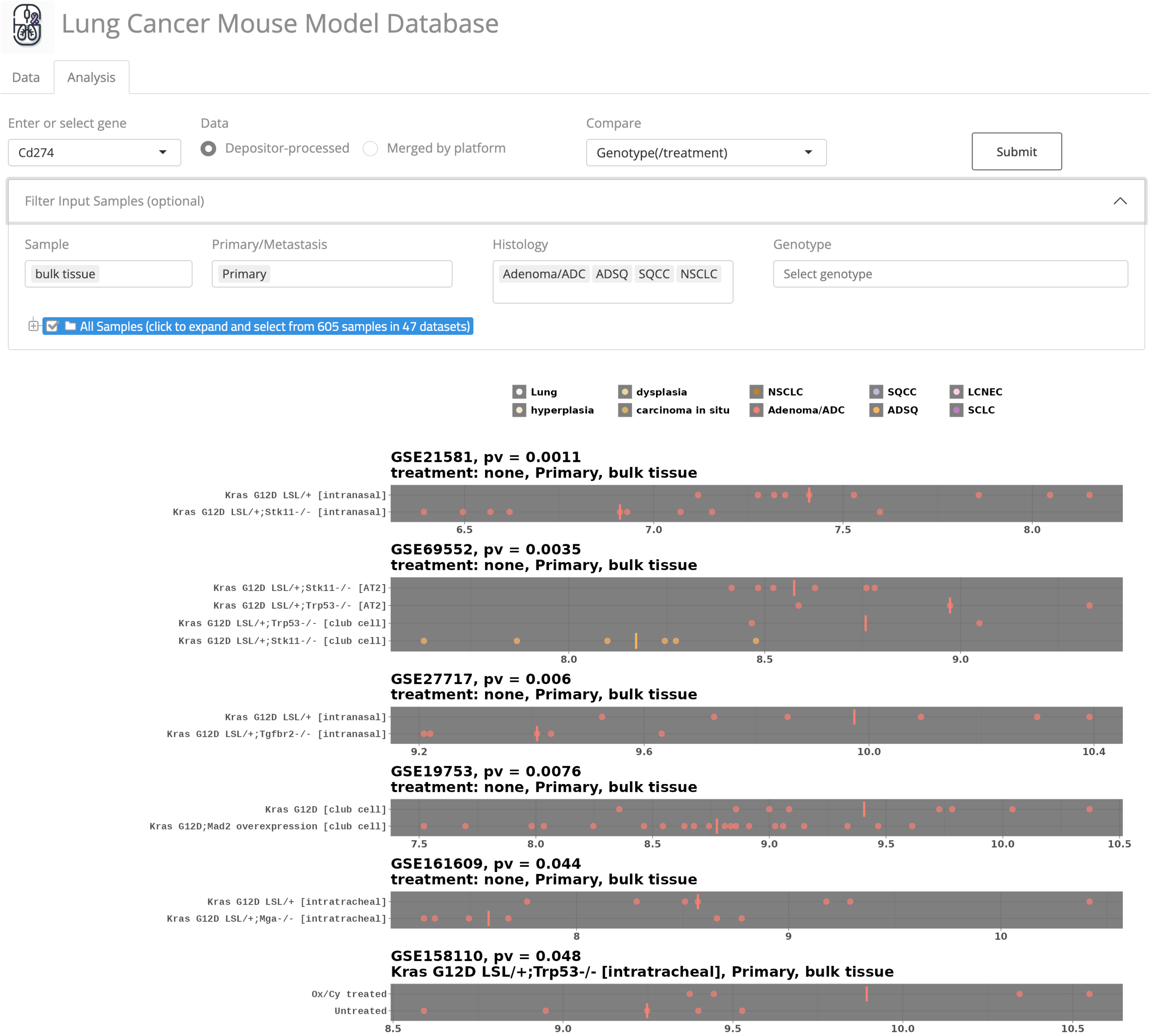
Interactive visualization of gene expression across multiple datasets. This figure features the web application’s capability for users to interrogate the expression of a selected gene, *Cd274* (PD-L1), across a range of datasets. The “Depositor-processed” option leverages the original data processed in the deposited datasets, optimizing the within-dataset comparisons. Users can tailor the analysis by applying filters via the dropdown menu. After selecting the appropriate parameters and clicking ‘Submit,’ the application generates dot plots arrayed by the statistical significance of their expression differences, as assessed by one-way ANOVA. Displayed here are the top 6 datasets from the full results, giving users a snapshot of the gene expression landscape within the application’s extensive repository. Bars in each plot denote the group median.

To enable more focused analyses on treatment/carcinogenesis response and cancer progression, we devised two additional comparison options for analyzing gene expression: one for treatment comparisons from ten studies and another for examining differences between primary tumors and metastatic lesions from five studies. The treatment comparison tool is showcased by analysis of the B cell marker *Cd19* to reveal distinct trends in tumor microenvironments (**Figure 5a**). We observed a pronounced increase in *Cd19* expression, indicative of B cell infiltration, in a Braf-driven GEMM under MAPK inhibitor treatment (GSE145152 dataset), which aligns with tumor regression [24]. Conversely, a significant decrease in *Cd19* expression was noted in samples with tumor progression, such as in Kras-driven GEMMs treated with antioxidants (GSE52594 dataset) [25]. *Cd19* also increased in Egfr-driven GEMMs subjected to a high-fat diet (GSE119649) and a Kras-driven GEMM under a high-caloric diet (GSE56260), in line with previous findings that obesity creates a more inflammatory tumor microenvironment in mouse [26], as well as observation in patients that high body mass index (BMI) is independently associated with overall survival benefit from immune checkpoint inhibitor therapy in advanced NSCLC [27].

**Figure 5:**
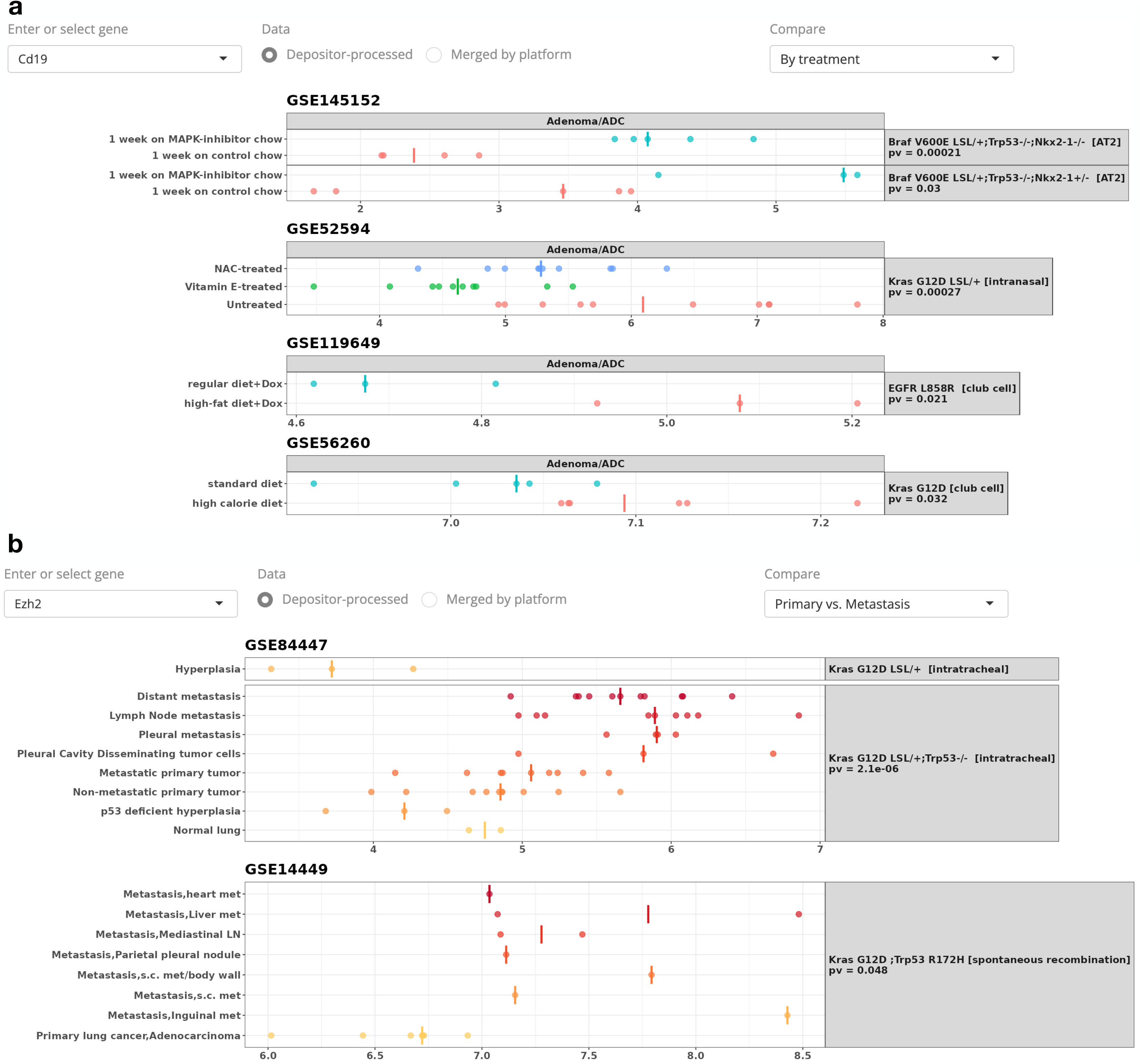
Gene expression comparison by treatment and primary/metastasis status. **a**. Expression of *Cd19* revealing B cell infiltration in various treatment contexts. Data points are categorized by treatment conditions under each genotype. **b**. Expression of Ezh2 in primary and metastatic tumor samples. Color gradient signifying the spectrum of metastatic progression stages. P-values from one-way ANOVA are indicated, and results were ordered by statistical significance. For conciseness, only the top 4 datasets out of 10 for *Cd19* (a) and the top 2 out of 5 for *Ezh2* (b) included in the snapshots. Bars in each plot denote group median.

The primary/metastasis comparison tool is exemplified by the examination of *Ezh2* expression, a component of the Polycomb Repressive Complex 2, which is implicated in gene silencing (**Figure 5b**). With the fine curation of metastatic status in samples from GSE84447 [28], we observe *Ezh2* expression increase with tumor invasiveness in Kras-driven models, which corroborates clinical findings that this chromatin modifier is associated with cancer progression and metastasis [29].

### Comparisons in merged reprocessed data

Analysis of reprocessed data merged by platform enables cross-study comparison. Users may select from six platforms with two or more merged datasets and further filter the input sample (as in **Figure S5**). We provide two visualization approaches for analyses. The first approach is to generate a dotplot with samples colored by histology and ordered by the median expression of the user-defined gene in groups stratified by a combination of data source, genotype, treatment, primary/metastasis status, and sample type. In the reprocessed RNA-seq data, this gives rise to 115 unique groups and creates a very extensive plot. We refined our selection to primary tumors from the RNA-seq reprocessed data and examined the expression of *Cd19*. The lowest expression is found in sorted cancer cells and samples with CD45 depletion (**Figure 6**, bottom), as expected from the depletion of immune cells. Bulk tissue samples with the lowest *Cd19* expression are from SCLC, consistent with the immune cold nature of this histological subtype [30-32]. The highest expression of *Cd19* is found in dysplasia samples derived following treatment with the alkylating agent N-nitroso-tris-chloroethylurea (NCTU), potentially due to abundant neoantigen resulting from carcinogen treatment (**Figure 6**, top). Users may select from additional profiling microarray platforms. For example, analysis of *Ezh2* expression in reprocessed data of Mouse430_2 reveals its expression is much higher in SCLC compared to NSCLC samples (**Figure S6**), as previously established [33, 34].

**Figure 6:**
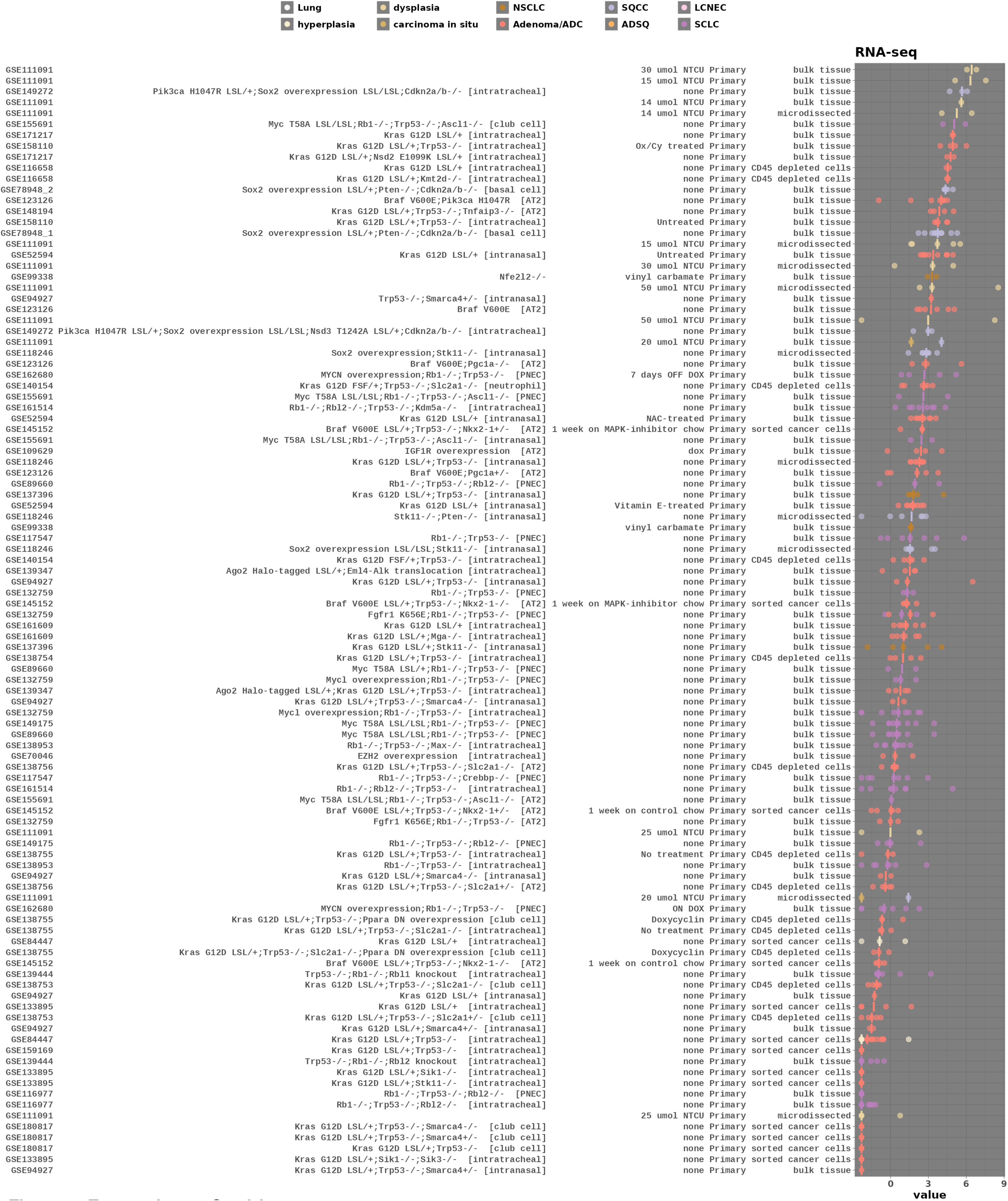
Expression of Cd19 in reprocessed RNA-seq data. Each dot represents a unique sample, colored according to histology, and ordered by the median expression of Cd19. The inputs are filtered to display primary tumors only. Median of the group is shown as a bar for each row.

The second visualization option generates a two-dimensional PCA plot, with sample points colored based on variables such as gene expression, histology, primary/metastasis status, sample type, or data source. In **Figure 7a**, we demonstrate this with RNA-seq samples colored to reflect *Ascl1* expression—a neuroendocrine lineage transcription factor instrumental in SCLC pathogenesis [35, 36]. Corroborating the histological segregation observed in **Figure 3e**, we found that samples characterized by lower PC1 scores—typical of SCLC—exhibited elevated *Ascl1* expression. Notably, some adenocarcinoma (ADC) samples, despite having higher PC1 scores, also showed high *Ascl1* levels. Our interactive plots equipped with informative tooltips reveal sample details, indicating that these outliers are ADCs from a model with constitutively-active *Fgfr1*^*K656E*^ in an *Rb1/Trp53*-deficient background [37], typically used to study classic SCLC (**Figure 3b**). While FGFR1 activation in this model has reduced *Ascl1* expression compared to classic SCLC tumors [37], the *Ascl1* levels are still higher than ADC tumors from other models (**Figure S7**).

**Figure 7:**
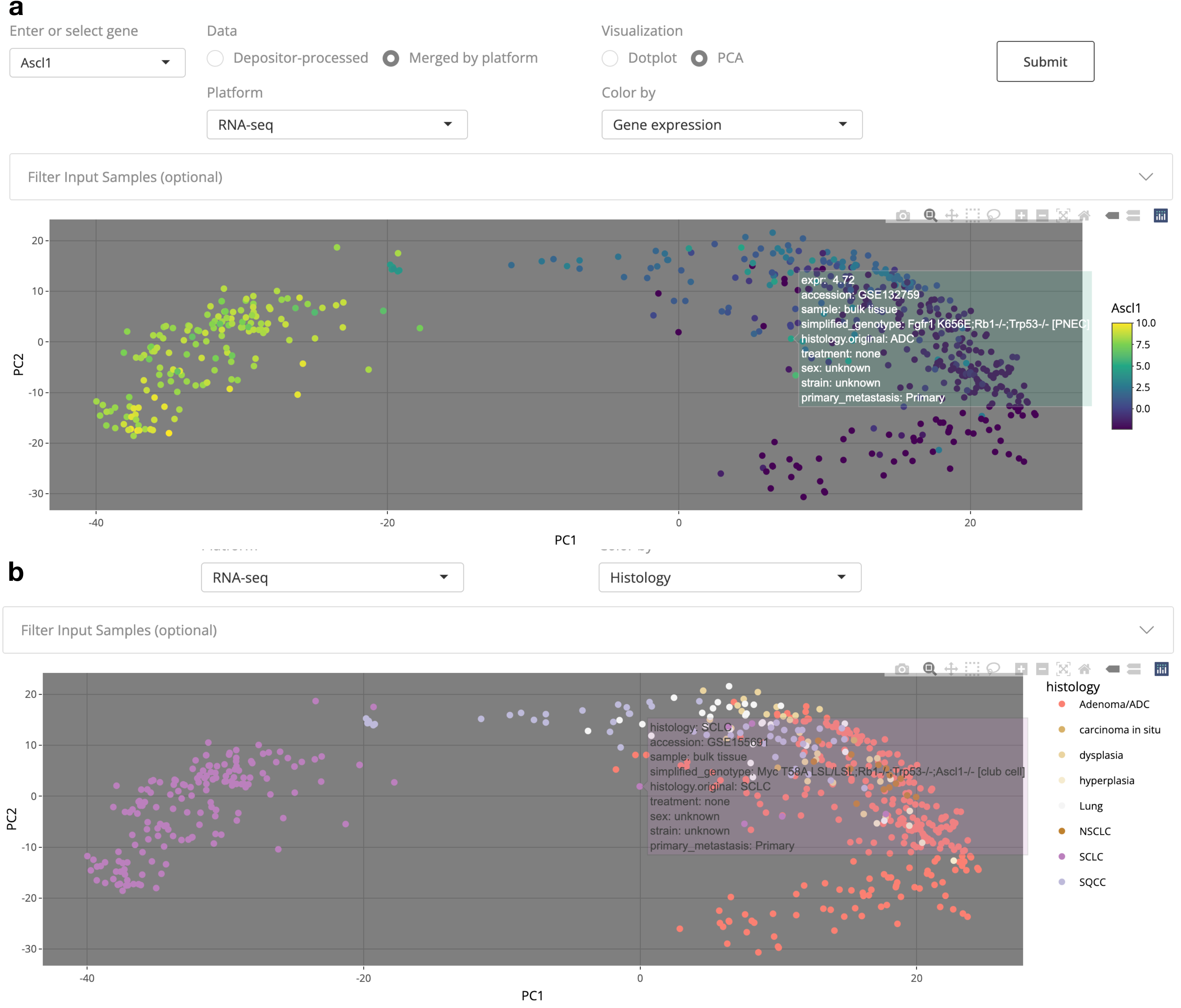
Interactive visualization of gene expression in reprocessed data merged by platform. **a**. PCA plot of reprocessed RNA-seq samples, color-coded by the expression of *Ascl1*, a neuroendocrine lineage transcription factor highly expressed in SCLC. The interactive tooltip uncovers the origin of an outlier sample with elevated Ascl1 levels as an ADC sample from a Rb1/p53 deficient model featuring Fgfr1 activation. **b**. PCA plot colored by histology. Details of a SCLC sample located near the NSCLC samples are read. This outlier sample has *Ascl1* knocked out, which explains the loss of neuroendocrine gene expression that renders the transcriptomic profile more similar to NSCLC.

In another example using the PCA plot with histology color mapping, there are a few notable outliers among the NSCLC samples that are identified as SCLC (**Figure 7b**). This particular discrepancy is clarified upon recognizing that these SCLC samples have undergone *Ascl1* knockout, leading to a complete loss of neuroendocrine cell fate [36] Consequently, ASCL1 knockout in a SCLC model makes the transcriptomic landscape of the SCLC sample more akin to that of NSCLC samples, explaining its outlier position in the PCA plot. Users also have the option to visualize data points according to the data source; the interactive plots enable users to selectively focus on, or exclude, samples from specific sources by clicking or double-clicking on dataset identifiers, thereby providing a clearer understanding of the underlying data distribution across studies (**Figure S8**).

## Discussion

Our LCMMDB presents a curated compendium of transcriptomic data covering 1,354 samples from 71 studies, summarizing a vast array of lung cancer mouse models. This resource interrogates the genetic aberration landscape across 859 GEMM tumors, providing an unprecedented platform for cross-study comparison. Our collaborative approach, engaging with data depositors, has ensured the integrity and enhancement of the database, leading to its current comprehensive state.

However, we need to consider the limitations inherent to the database’s scope. The LCMMDB is founded on transcriptomic data, which excludes mouse models lacking such characterization. This limitation underscores the need for an inclusive approach that considers unpublished or less-publicized models to achieve a comprehensive and representative overview of genetic alterations in lung cancer. The dynamic nature of scientific research also necessitates the LCMMDB to be a living database, with ongoing updates and expansions informed by both community feedback and continual data discovery. Future versions will integrate additional datasets, reflecting the latest advancements and filling in gaps identified through collaborative suggestions and our active searches. The candidate datasets to be included in the next update are listed in **Supplementary Table S6**. These include datasets from a more recent GEO/ArrayExpress screen and datasets suggested by the community, such as toxicology studies of mouse lungs treated with carcinogens. We will also continue to develop the collaborator login system, enabling researchers to privately assess their data alongside public datasets. While not expounded upon in this manuscript, this function highlights the platform’s potential for fostering collaborative research endeavors.

It is important to note the caution required in interpreting the reprocessed data. While standardization efforts have been rigorous, batch effects from diverse experimental and genetic backgrounds may still be present. Future updates will aim to support meta-analytical capabilities and provide insights from comprehensive cross-transcriptomic evaluations. Central to the LCMMDB’s utility is its facilitation of comprehensive comparisons between mouse models, additional preclinical model data, such as patient-derived cell lines, patient-derived xenografts, syngeneic mouse models, and human lung cancer data. This alignment is crucial for translating preclinical findings to clinical relevance, aiding in the development of personalized therapies. The database’s current iteration lays the groundwork for such comparative studies, which we plan to explore in-depth in subsequent analyses.

In sum, the LCMMDB offers a robust framework for the exploration of gene expression data within mouse models, setting the stage for additional comprehensive analyses that have the potential to unveil new discoveries and guide the design of future models for a more accurate reflection of human lung cancer.

## Acknowledgments

This study is supported by funding from UTSW ACS-IRG (IRG-21-142-16), P50CA70907, U24CA213274, R01GM140012, R01GM141519, R01DE030656, R01CA244841 (TGO), U01AI169298, U01CA249245, U01AI156189, R35CA22044901, U01CA213338, R35GM136375, and R35CA263816. The authors declare no competing interests.

L.C. designed the study, performed the analyses, and wrote the manuscript. Y.G. reviewed additional datasets for inclusion in a future update. T.G.O. and J.D.M. provided critical inputs for the manuscript. T.G.O., R.J.D., J.D.M., and Y.G. helped edit the manuscript. L.C., R.J.D., G.X., T.G.O., C.M.R., and Y.X. obtained funding to support this work. We thank the lung cancer mouse model community for their efforts in helping to curate this resource. Among the authors, G.A., V.A., J.E.B., S.B., J.B., T.C., C.P.C., B.M.G., D.J., R.J., J.M.K., M.L. (Lee), P.L., Y.L., M.L. (Lopez), R.M. (Martinelli), P.K.M., S.A.M., S.M., H.M., R.M. (Moorehead), E.E.M., S.N., T.G.O.,

M.G.O., A.R.P., C.A.P., G.R., M.S., E.S., R.S., K.S., T.T., K.T., Y.X., E.V., and M.W. provided dataset confirmation, corrected or improved data curation, and feedback; M.G.O., M.W., T.G.O., and J.D.M. suggested additional datasets to include. In addition to the listed authors, we want to acknowledge additional data curation confirmation/correction help from Drs. Andrea Ventura, Anneleen Daemen, Anton Berns, Bob Stearman, Casanova Emilio, Celeste Simon, Cong Yan, Emmy Verschuren, Francesco Demayo, Julien Sage, Karsta Luettich, Kwok-Kin Wong, Meylan Etienne, Shivani Srivastava, and Thomas Russell; and dataset suggestions from Drs. Karsta Luettich, Julien Sage, and Thomas Russell.

