## Supplementary Figures 1-8 for "A Lung Cancer Mouse Model Database"

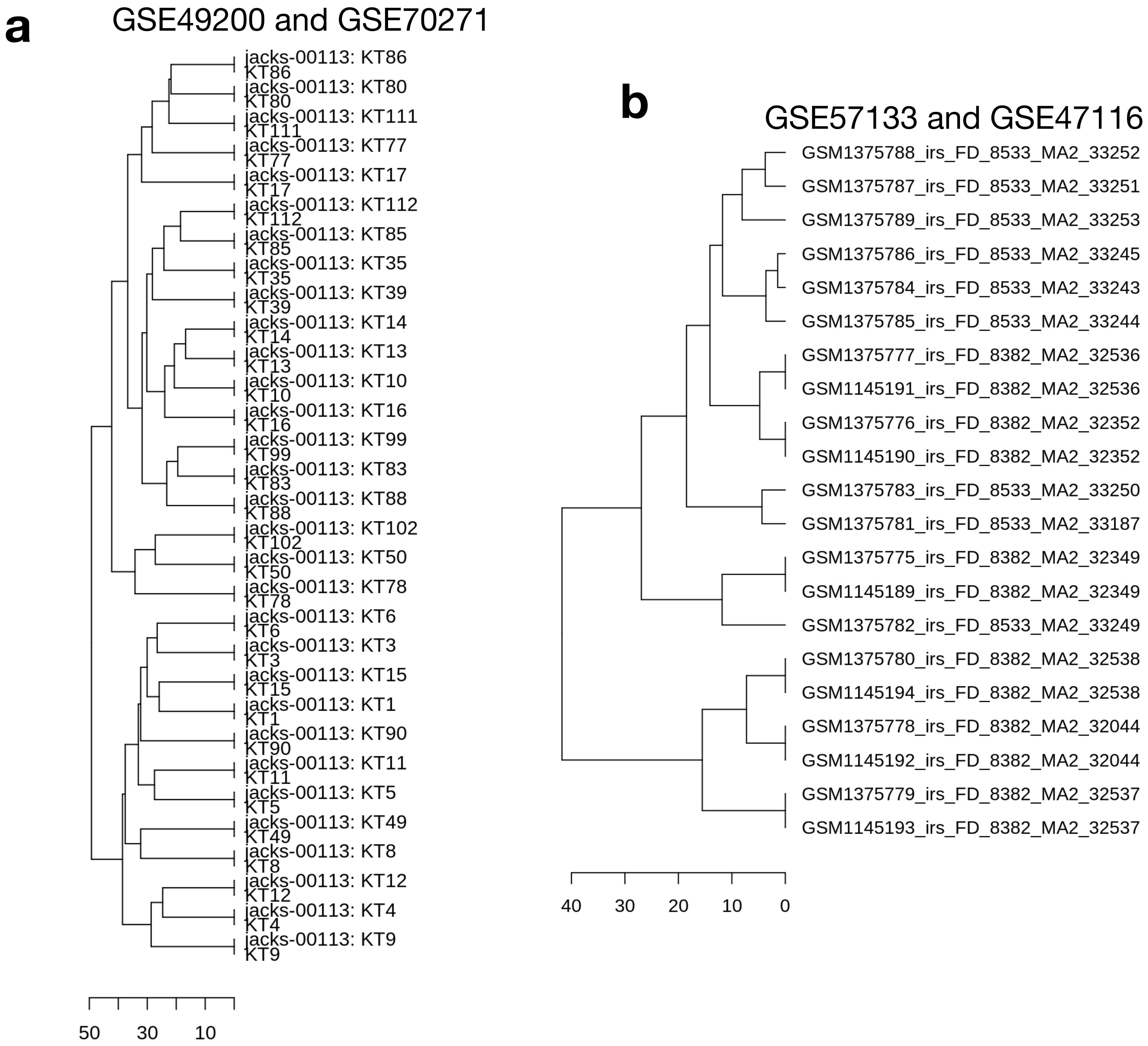


**Figure S1. Identification of data redundancy by hierarchical clustering.**
**a**. All 50 samples from GSE49200 overlap with 50 of the 58 samples from GSE70271 (only the overlapping samples are included). Data were reprocessed with MG-U74 probe anntation files. **b**. All 6 samples from GSE47116 overlap with 6 of 10 samples from GSE57133. Data were reprocessed with the Mouse430_2 probe annotation files. Top 10 principal components from the 1000 most variable genes were used to compute Euclidean distance for hierarchcal clustering.

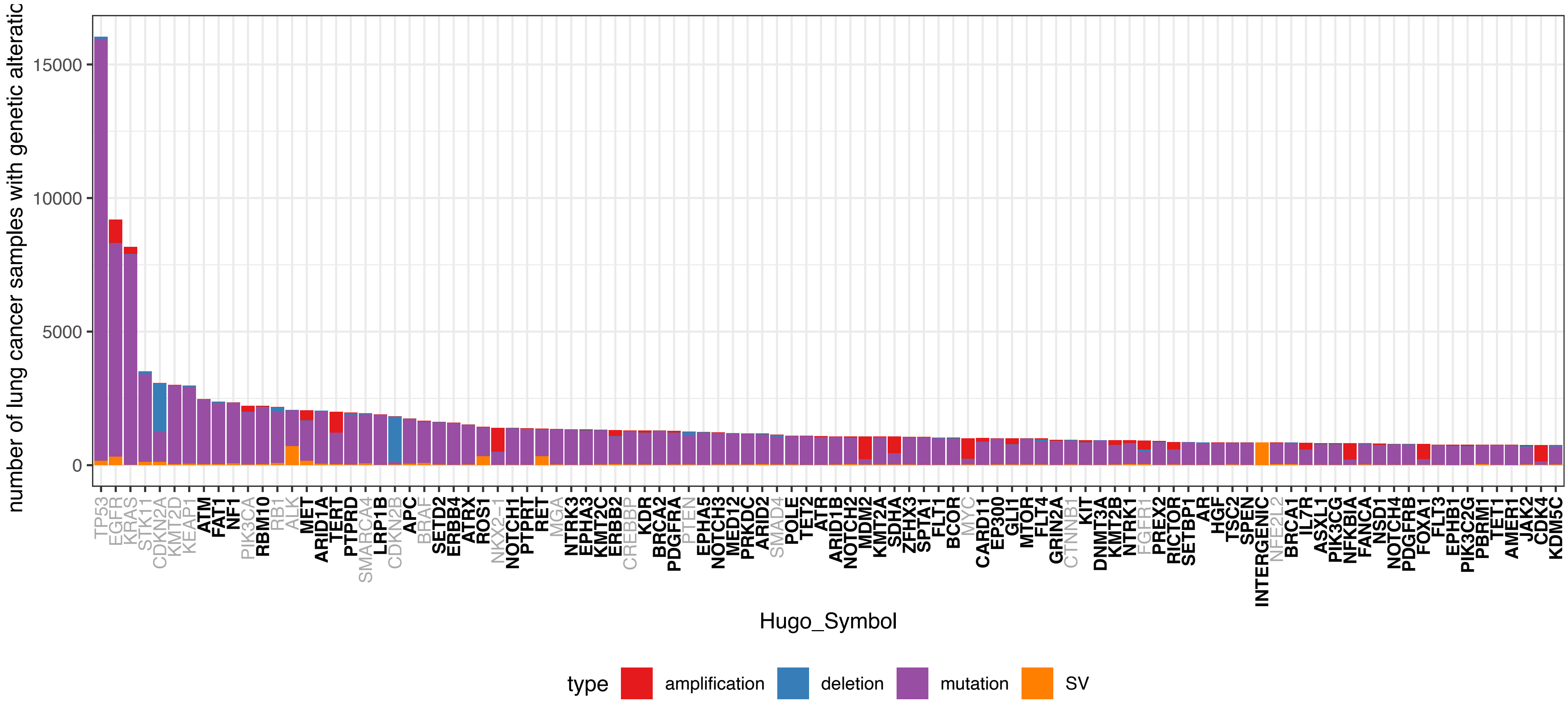


**Figure S2. Top 100 frequent genetic mutations in realworld lung cancer patients**
This bar chart showcases the top 100 most frequent genetic mutations observed in lung cancer patients, derived from the analysis of AACR GENIE v15 data, encompassing 29,701 samples from 25,683 patients. It includes panel sequencing mutations, copy number alterations, and large-scale chromosomal rearrangements. The y-axis indicates the number of samples with each type of genetic alteration, and the x-axis lists the genes in descending order of alteration frequency. Genes that are absent in the GEMM tumors from the LCMMDB are distinguished by bold and darkened labels on the x-axis.

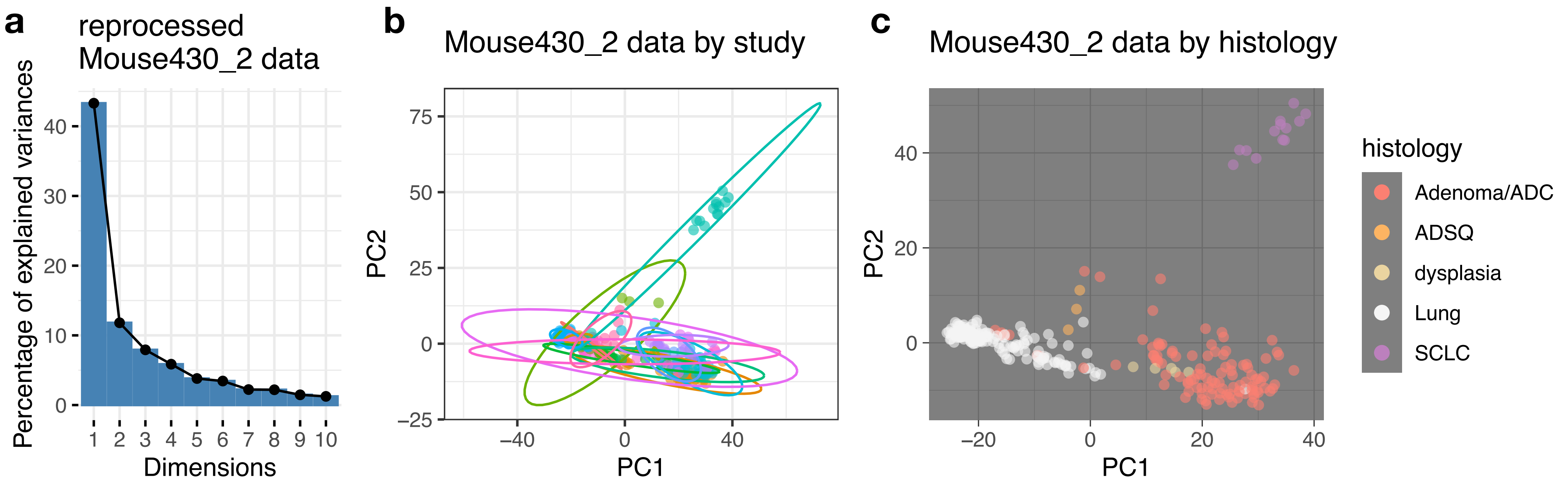


**Figure S3. PCA of reprocessed Mouse430_2 microarray data**

**a**. Variance explained by the top ten principal components in the PCA of the reprocessed Mouse430_2 data, highlighting the significant proportion of variance captured by the first two dimensions. **b**. PCA biplot of Mouse430_2 data by study, with ellipses representing the dispersion of samples within each dataset. Each color and ellipse corresponds to a unique study, illustrating the variability and overlap among studies. **c**. PCA scatterplot of Mouse430_2 data by histology type, showing the distribution of samples across the first two principal components. Note that the samples reprocessed for Mouse430_2 do not contain metastasis samples so separation by primary/metastasis status is not visualized here.

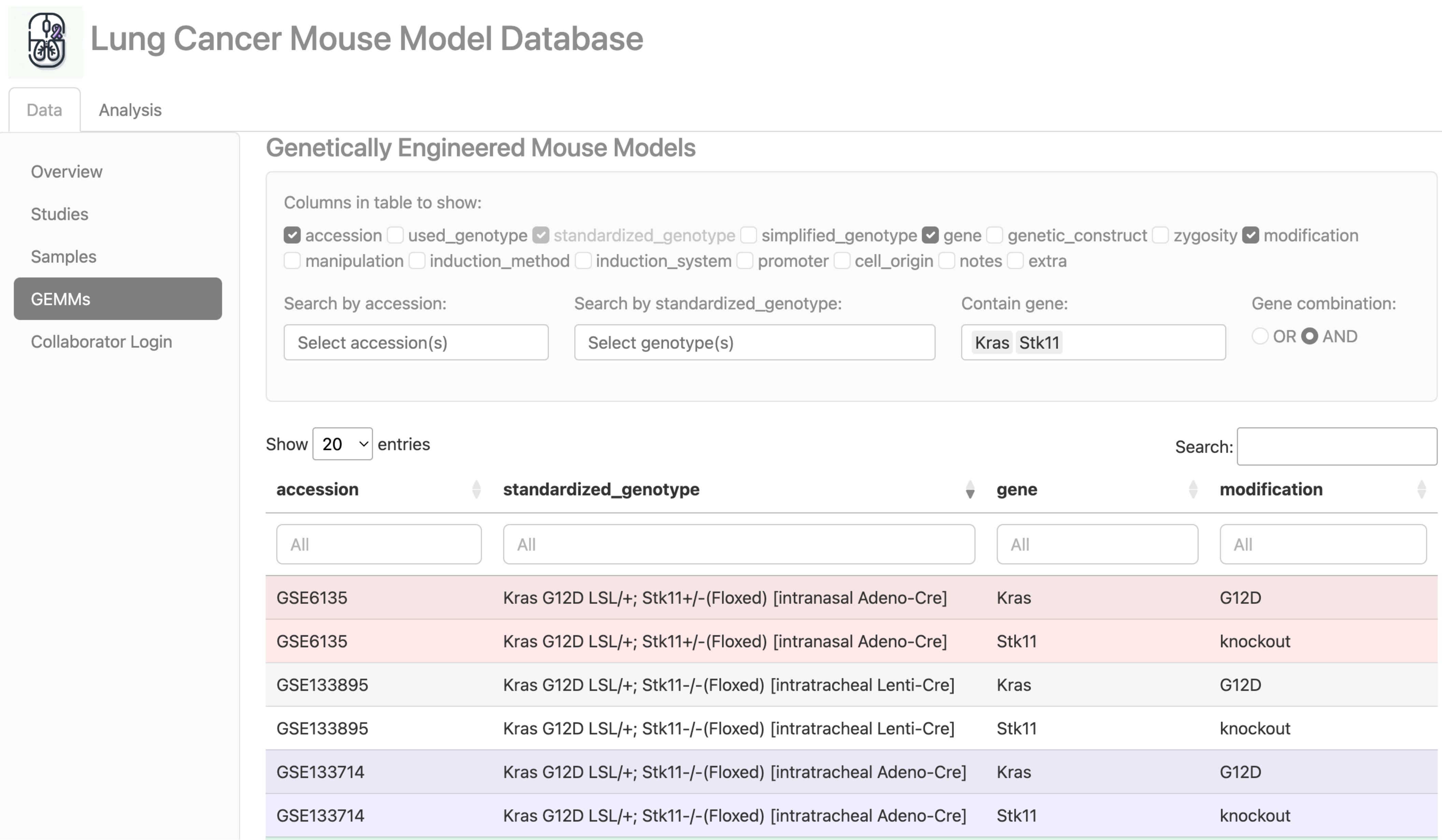


**Figure S4: User interface of the GEMMs table in the LCMMDB web application.**
This screenshot showcases the GEMMs table interface within the LCMMDB web application. The table allows users to view and filter genetic alteration data, displaying one gene per row. Each row is color-coded to correspond to a unique genotype within a study, enhancing visual differentiation. The interface provides options for users to display selected columns, and features a filtering system for searching specific gene combinations or alterations. This functionality aids in the efficient curation and analysis of complex GEMM design.

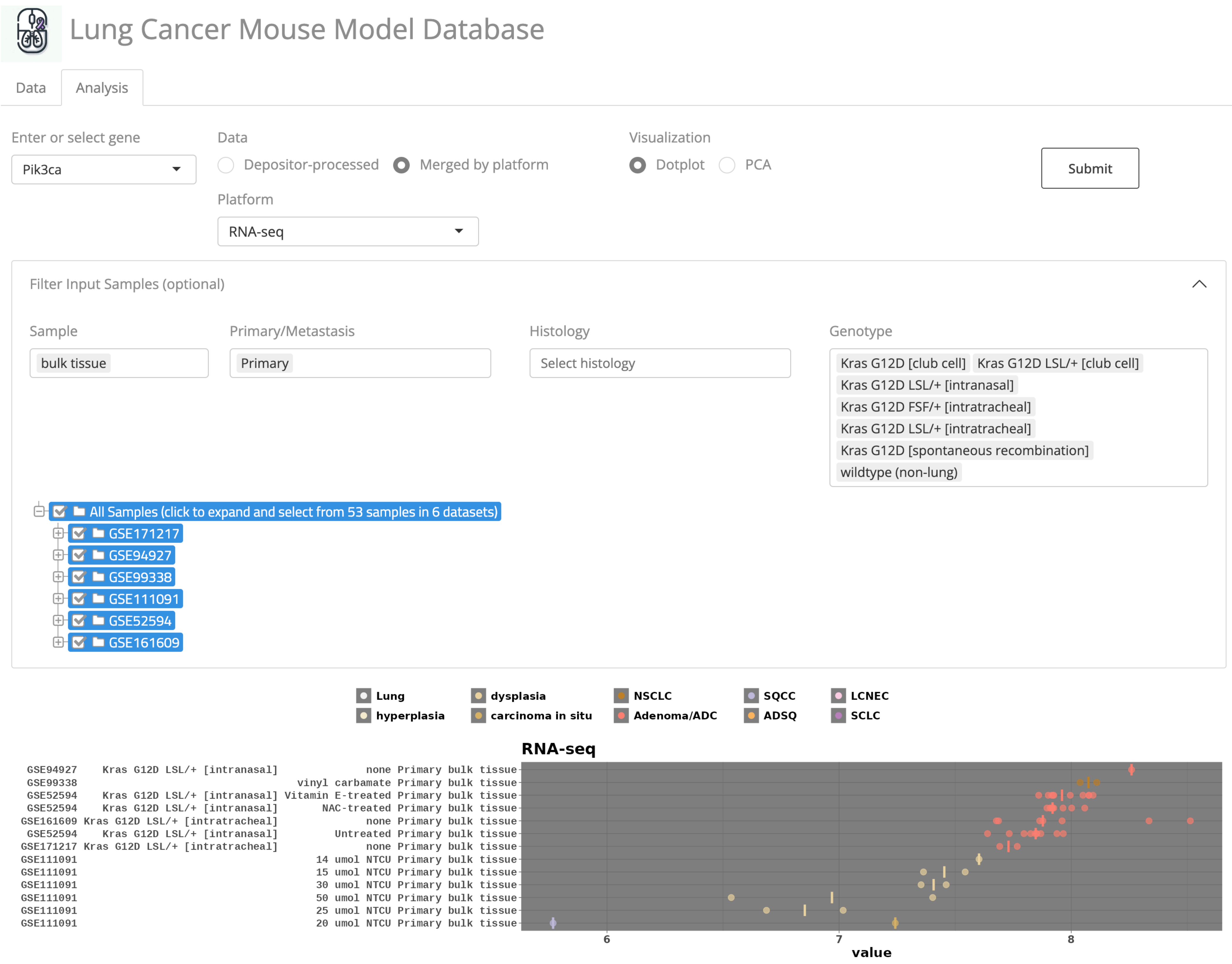


**Figure S5: Custom sample filtering and selection**
This figure demonstrates the web application's capability for tailored sample filtering, for generating dot plots from depositor-processed or uniformly reprocessed datasets. Within the expandable menu, users can specify sample characteristics such as type, primary versus metastatic status, histology, and genotype. The sample set is further navigable via a dynamic tree structure categorized by data source, which adapts in real time to platform selection and applied filters for sample attributes. Here, we illustrate the selection process for primary lesion bulk tissue samples with Kras G12D or wildtype genotypes. The subsequent dot plot visualization, focusing on the Pik3ca expression profile within RNA-seq data, reveals pronounced expression in GEMM tumors relative to wildtype mice with carcinogen-induced lesions.

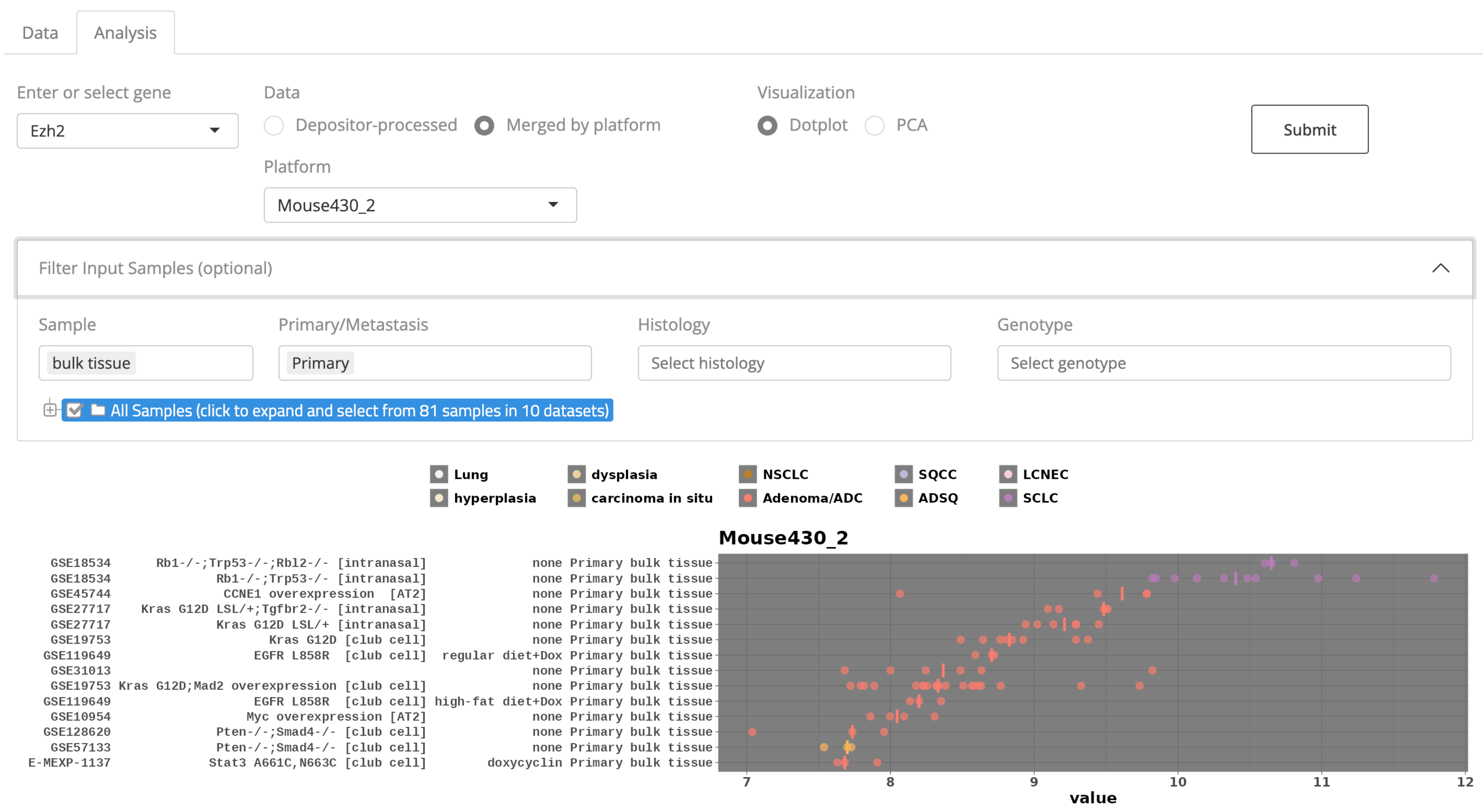


**Figure S6. Dotplot of Ezh2 expression across various samples profiled by the Mouse430_2 microarray platform.** Each dot represents a unique sample, colored according to histology, and ordered by the median expression of Ezh2. The plot is refined to display bulk tissue primary tumors, highlighting the differential expression levels between small cell lung cancer (SCLC) and non-small cell lung cancer (NSCLC), with a notable elevation in SCLC samples.

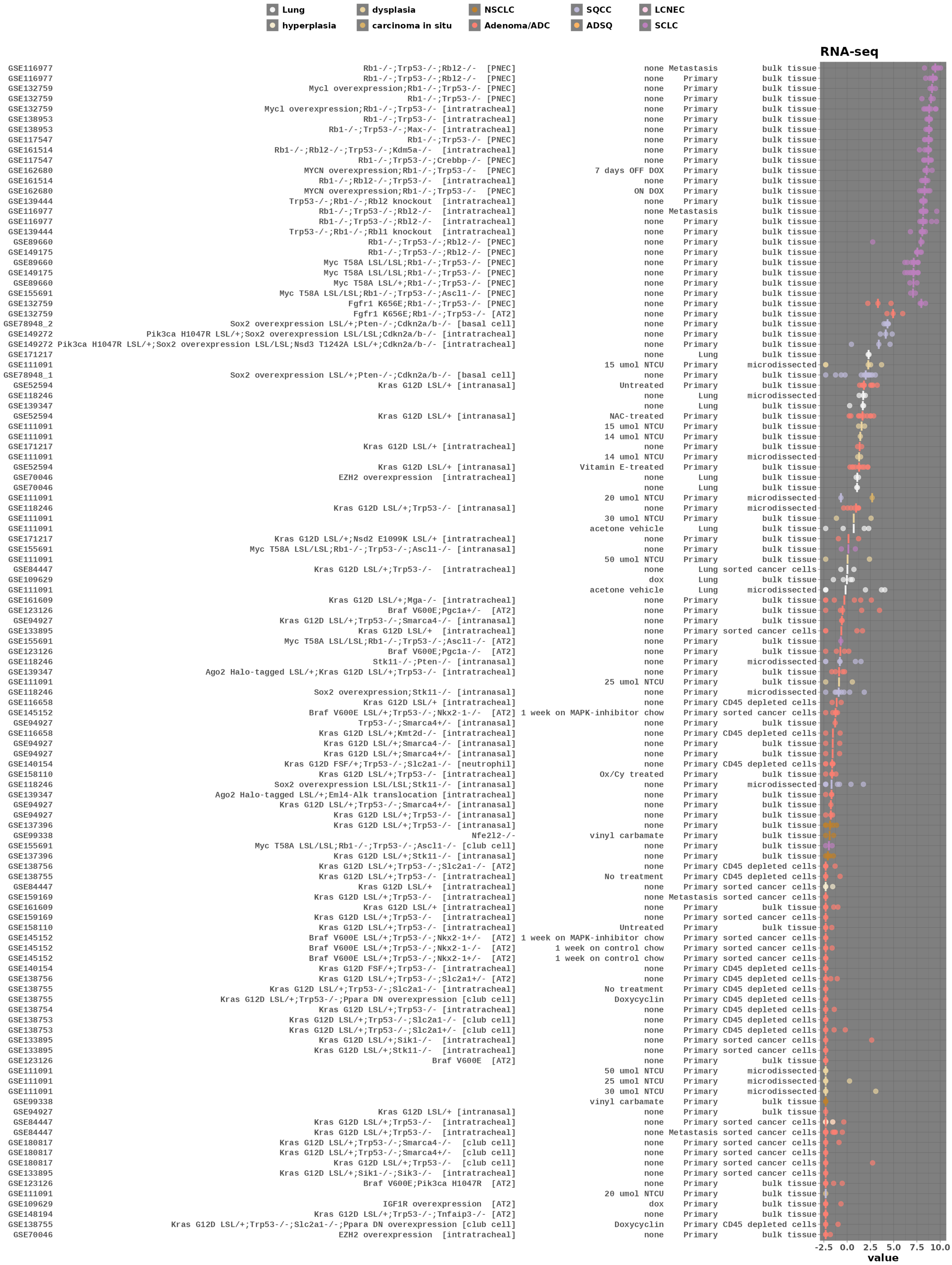


**Figure S7. Ascl1 expression across reprocessed RNA-seq datasets**

Note that in GSE132759, despite ADC histology in tumors with Fgfr1 mutationin Rb1/Trp53 loss background, remnant Ascl1 expression was detected.

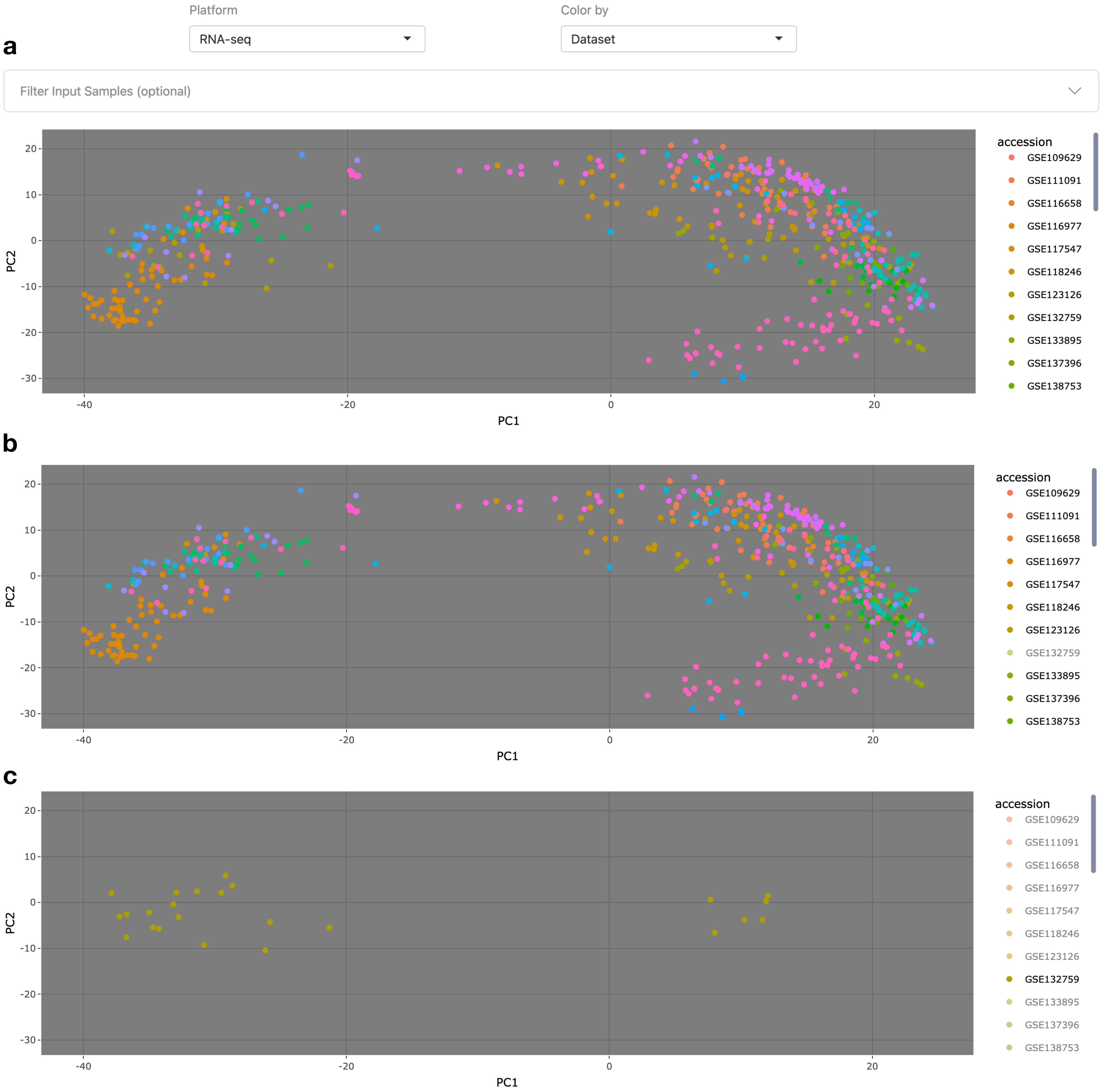


**Figure S8. Samples colored by data source.**
**a.** Samples colored by data source to illustrate diversity in reprocessed data by platform. **b.** Single click on “GSE132759” hides data points under this accession id. **c.** Double click on “GSE132759” hides all other datapoints except for those under this accession id.
